# Impaired BDNF-TrkB trafficking and signalling in Down syndrome basal forebrain neurons

**DOI:** 10.1101/2025.07.15.664857

**Authors:** Emily Blackburn, Nicol Birsa, Oscar M. Lazo, Elizabeth M.C. Fisher, Giampietro Schiavo

## Abstract

Brain derived neurotrophic factor (BDNF) and its receptor tropomyosin-related kinase B (TrkB) play crucial roles in neuronal development, synaptic transmission, and neuroplasticity. Deficits in BDNF/TrkB signalling and trafficking have been identified in several neurodegenerative diseases, including Alzheimer’s disease (AD). Individuals with Down syndrome (DS) are at an increased risk of developing AD compared to the general population. Basal forebrain neurons (BFNs) are among the first to degenerate in AD and DS, but the mechanisms underlying their vulnerability remain unclear. Using BFNs derived from the Dp1Tyb mouse model of DS, we investigated neurotrophic signalling and trafficking deficits in AD-DS. We found enlarged early endosomes and elevated levels of active Rab5, a GTPase critical for early endosome formation, in Dp1Tyb BFNs. These abnormalities were associated with impaired transport of internalised TrkB from axon terminals to the soma. Using microfluidic devices, we demonstrated that axonal BDNF stimulation enhanced signalling endosome dynamics in wild-type but not Dp1Tyb BFNs, which is likely due to impaired axonal ERK1/2 signalling. Our findings establish a link between Rab5 hyperactivation, endosomal dysfunction, and impaired ERK1/2 signalling, highlighting the interplay between trafficking and neurotrophic signalling, and underscore the importance of targeting endolysosomal and signalling pathways to mitigate neuronal dysfunction in AD-DS.

## Introduction

Neurotrophic signalling is crucial for the differentiation, growth and survival of neurons, as well as synaptic transmission and neuroplasticity. Neurotrophins, namely nerve growth factor (NGF), brain-derived neurotrophic factor (BDNF), neurotrophic factor-3 (NT-3) and neurotrophic factor 4 (NT-4), are a family of structurally related proteins that bind with high affinity to specific tyrosine kinase receptors (TrkA-C) (1). Following expression and secretion from target tissues, neurotrophins bind to Trk receptors mainly at axon terminals, activating the phosphoinositide 3-kinase (PI3K)-Akt, mitogen-activated protein kinase (MAPK-ERK) and phospholipase C-gamma (PLCγ) signalling pathways (1). All four neurotrophins also bind the p75 neurotrophin receptor (p75^NTR^), which interacts with Trk receptors, modulating their function (2).

Neurotrophic signalling promotes both local and long-distance effects; at nerve terminals, it enables axonal branching and elongation, as well as synapse maturation and plasticity (3–5). Distal action requires the propagation of signals regulating gene expression, survival, development and dendritic arborisation to the cell body via the axonal retrograde transport of activated neurotrophin-Trk complexes (5–8). This transport is mediated by the recruitment of cytoplasmic dynein onto signalling endosomes (9–11), highlighting a critical relationship between axonal retrograde transport and long-distance propagation of neurotrophic signals. The internalisation of activated neurotrophin-Trk complexes from the plasma membrane, and the subsequent formation and maturation of these organelles, is regulated by Rab proteins - monomeric GTPases that cycle between an active, GTP-bound, membrane-associated state and an inactive GDP-bound cytosolic form (12).

Following internalisation, activated neurotrophin-Trk complexes are targeted to early endosomes which are regulated by Rab5, a GTPase crucial for receptor sorting and homotypic fusion of these organelles, among other functions (13). The complex is then returned to the plasma membrane via Rab4/11 recycling endosomes or sorted to Rab7-positive late endosomes for delivery to the soma, or to lysosomes for degradation (14, 15). In primary neurons, following activation by BDNF, TrkB has been shown to localize to both Rab5 and Rab11 positive compartments in dendrites and the cell body, a process crucial for dendritic branching (16, 17). Furthermore, ERK1/2-dependent CREB phosphorylation in the nucleus also depends on both Rab5 and Rab11 (18) as well as Rab10 (19). Upon BDNF treatment, Rab10-positive compartments are mobilised towards axon terminals, facilitating the sorting of TrkB into the axonal retrograde transport pathway (19). Thus, the precise spatial and temporal association of Trk receptors with specific Rab membrane compartments is crucial for the coordination between local effects and long-distance propagation of neurotrophic signalling in neurons.

Deficits in signalling and trafficking of neurotrophic receptors have been identified in many neurodegenerative diseases (20, 21). For example, disrupted axonal transport is an early event in Alzheimer’s disease (AD), occurring prior to the formation of Aβ plaques and neurofibrillary tangles in familial AD mouse models (22–24). Trisomy of chromosome 21 (Chr21), which gives rise to Down syndrome (DS), is the most common genetic cause of AD (referred to as AD-DS), with most individuals having AD-like neuropathology by the age of 40 (25). The key dosage-sensitive Chr21 gene underpinning the increased risk of AD is the *APP* gene, which encodes the amyloid precursor protein; however, several other proteins encoded by genes on Chr21 have also been suggested to play a role in AD-DS (26). Notably, the cholinergic system is particularly vulnerable in both DS and AD, as cholinergic basal forebrain neurons (BFNs) rely heavily on the retrograde transport of neurotrophins to maintain their cholinergic identity (27, 28).

Several studies have found abnormalities in neurotrophin signalling in AD and AD-DS, including reduced neurotrophin and Trk receptor levels (29, 30), impaired neurotrophin transport (31–34) and deficits in BDNF-related signalling via ERK and AKT (34). Furthermore, impaired endosome morphology and function are early events in AD and DS human *post-mortem* tissue and mouse models (34–44). It has been suggested that APP, and its associated ß-secretase cleaved, membrane-bound carboxy terminal fragment (ß-CTF), induce this pathological phenotype via Rab5 hyperactivation (39, 42, 44–46). Additionally, enlarged endosomes convey trophic signals less effectively both *in vitro* and *in vivo* (32, 43). Thus, the available evidence suggests that defects in neurotrophic signalling are not only directly related to neurotrophin/Trk levels but also implicate impaired axonal retrograde trafficking of signalling endosomes in AD and DS.

Here, we use an established model of DS, the Dp1Tyb mouse, which has an extra copy of the human Chr21 (Hsa21) orthologous genes on mouse chromosome 16 (Mmu16), comprising 63% of all Hsa21-orthologous mouse genes (47, 48). However, most research exploring the role of early endosomes and neurotrophic signalling in the pathogenesis of AD-DS has been conducted using the Ts65Dn mouse. This mouse carries 60 annotated genes that are not homologous to Hsa21. Included in this group are at least four genes (*Snx9*, *Synj2*, *Dynlt1*, *Sylt3*) encoding proteins involved in the endocytic pathway and/or vesicular trafficking (49). Indeed, many of these genes could contribute to Ts65Dn phenotypes and act as confounding factors in the analysis of the pathomechanism of AD-DS.

The aim of this work was to address whether BFNs, a highly vulnerable neuronal population in DS and in AD, isolated from Dp1Tyb embryos display neurotrophic signalling and trafficking deficits. We confirmed the presence of enlarged early endosomes in BFNs and identified increased levels of active Rab5 specifically in Dp1Tyb neurons. Furthermore, we found that BDNF enhances the transport dynamics of BDNF/TrkB signalling endosomes in WT but not Dp1Tyb BFNs, which is likely due to impaired ERK1/2 signalling in Dp1Tyb BFN axons.

## Results

### Early endosome morphology and function are altered in Dp1Tyb basal forebrain neurons

To identify whether BFNs derived from Dp1Tyb mice display early endosome abnormalities, we stained DIV14 neurons for EEA1, an established early endosomal marker (**Figure 1A**). Dp1Tyb BFNs displayed significantly enlarged early endosomes, >50% larger than that seen in WT BFNs (**Figure 1B**). Furthermore, there was a ∼60% increase in the number of larger endosomes and a ∼13% reduction in the number of smaller endosomes in Dp1Tyb neurons (**Figure 1C**). Early endosomes were also more numerous in Dp1Tyb BFNs (**Figure 1D**).

**Figure 1:**
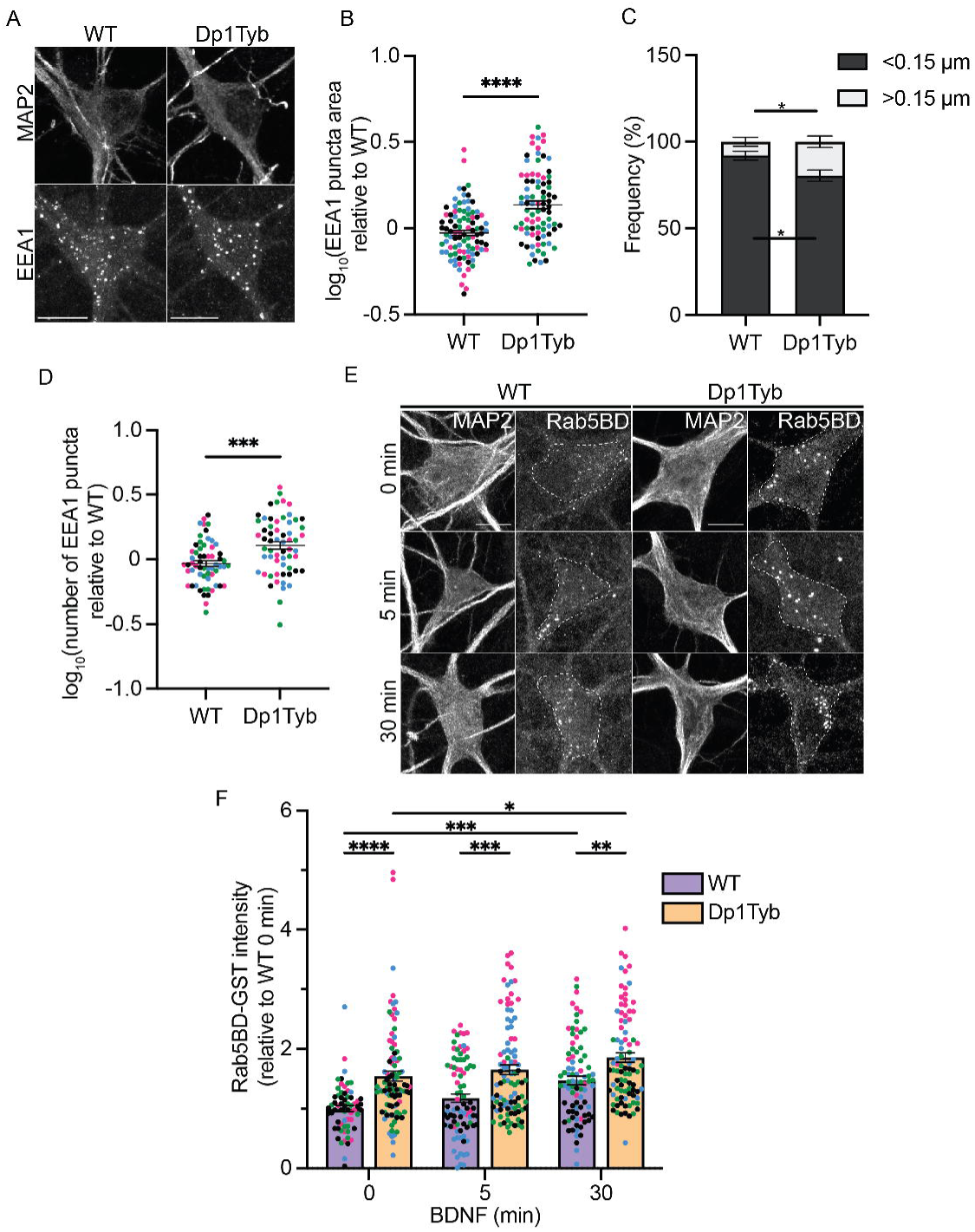
Early endosomes are enlarged and display increased Rab5 activity in basal forebrain neurons isolated from Dp1Tyb embryos. (A) Representative confocal images of EEA1 immunofluorescence in WT and Dp1Tyb BFNs at DIV14. (B) EEA1 positive puncta were significantly larger in Dp1Tyb neurons compared to WT (Student’s *t* test; ****, p <0.0001). (C) The size distribution of EEA1 positive puncta in Dp1Tyb BFNs showed a significant shift from smaller to larger binned areas compared to WT littermates (Student’s *t* test; *, p=0.0320). (D) Quantification of EEA1 puncta number in Dp1Tyb BFNs compared to WT (Student’s t-test; ***, p=0.0002). (E) Representative images of WT and Dp1Tyb neurons incubated with GST-Rab5BD to measure levels of GTP-Rab5 in the cell bodies (dashed lines) following 0, 5, 30 min BDNF treatment. (F) BDNF significantly increased GTP bound Rab5 in BFNs after 30 min treatment. GTP-Rab5 levels were significantly elevated in Dp1Tyb BFNs compared to WT at baseline and with BDNF treatment. (2-way ANOVA with Tukey’s multiple comparisons; BDNF; ****, p<0.0001, genotype; ****, p <0.0001. Multiple comparisons; ****, p <0.0001, ***, p=0.0009, **, p=0.0072, *, p=0.0368). Different coloured points represent cells from separate *N*s. Bars represent mean ± SEM. *N* = 4 biological replicates. 15-25 cells were imaged per *N*. Scale bar = 10 µm.

It has been suggested that Rab5 hyperactivation underpins the early endosomal enlargement in AD and AD-DS (50). As such, we wanted to ascertain whether there were increased levels of active Rab5 in Dp1Tyb BFNs. We used an *in-situ* approach using GST-fused to the Rab5 binding domain of Rabaptin-5 (Rab5BD) as a probe to specifically measure GTP-bound endogenous Rab5 (Rab5-GTP), followed by an immunostaining against GST (**Figure 1E**). Rabaptin5 is a Rab5 effector that preferentially binds active Rab5 (51). Prior to incubating with the Rab5BD probe, BFNs were treated with BDNF for 0, 5 or 30 min to determine whether BDNF stimulation induces an increase in Rab5 activity, as has been previously shown in primary hippocampal neurons (17).

We found that in WT neurons there was a significant increase in Rab5-GTP after 30 min BDNF treatment, confirming that BDNF increases Rab5 activity. Consistent with the increased size of early endosomes, Dp1Tyb BFNs already showed significantly higher levels of Rab5-GTP at baseline (0 min); these are comparable to those found in the stimulated WT neurons, indicating increased basal levels of active Rab5 (**Figure 1F)**.

As positive and negative controls for these experiments, we incubated SH-SY5Y neuroblastoma cells transfected with constitutively active (Rab5CA) or dominant-negative Rab5 (Rab5DN) mutants, with the GST-Rab5BD probe (**Supplementary** Figure 1A**, C**). Cells were also incubated with GST alone to rule out non-specific GST binding (**Supplementary** Figure 1B). Early biochemical studies indicate that the N133I mutant acts as a dominant-negative through poor prenylation and membrane binding (52). Thus, whilst it is expected to bind GST-Rab5BD, the lack of a punctate binding pattern further indicates the specificity of our assay.

Altogether, these results highlight that Dp1Tyb BFNs exhibit elevated baseline Rab5-GTP levels resembling WT neurons treated with BDNF, indicating an inherent increase in Rab5 activity in these neurons compared to WT controls.

### The BDNF-dependent enhancement of axonal transport of signalling endosomes is impaired in Dp1Tyb BFNs

Next, we wanted to investigate whether increased levels of active Rab5 and enlargement of early endosomes impacted the trafficking of BDNF-TrkB complexes. To test this hypothesis, we first determined whether WT and Dp1Tyb BFNs display differences in TrkB levels. As shown in **Supplementary Figure 2A, B**, although the expression of TrkB is developmentally regulated in BFNs, we found no differences between the two genotypes at each timepoint. While effective for detecting total TrkB expression, immunofluorescence quantification does not distinguish between the isoforms of this receptor. Thus, we used western blotting to differentiate between full length TrkB (TrkB-FL) and its truncated T1 isoform (TrkB-T1); however, our analysis revealed no significant difference in TrkB-FL, TrkB-T1 levels, or the FL/T1 ratio (**Supplementary Figure 2C, D).**

Next, to evaluate the axonal retrograde transport of BDNF-TrkB complexes in live neurons, we exploited the non-toxic binding fragment of tetanus neurotoxin bound to AlexaFluor-555 (H_C_T-555). H_C_T has been shown to share axonal retrograde transport carriers with neurotrophins and their receptors and is a reliable proxy of BDNF-TrkB signalling endosome dynamics (13, 53, 54). To facilitate axonal transport analysis, primary BFNs were grown in three-chamber microfluidic chambers (MFCs; **Figure 2A**). These devices allow cellular and fluidic isolation of axon terminals, whilst also reducing variability between different dishes through the projection of axons from the same pool of neurons into two fluidically-isolated compartments. DIV14 BFNs were starved of growth factors and axonal compartments were then supplemented with H_c_T-555 in the presence or absence of 50 ng/mL BDNF 30 min prior to imaging and tracking.

**Figure 2:**
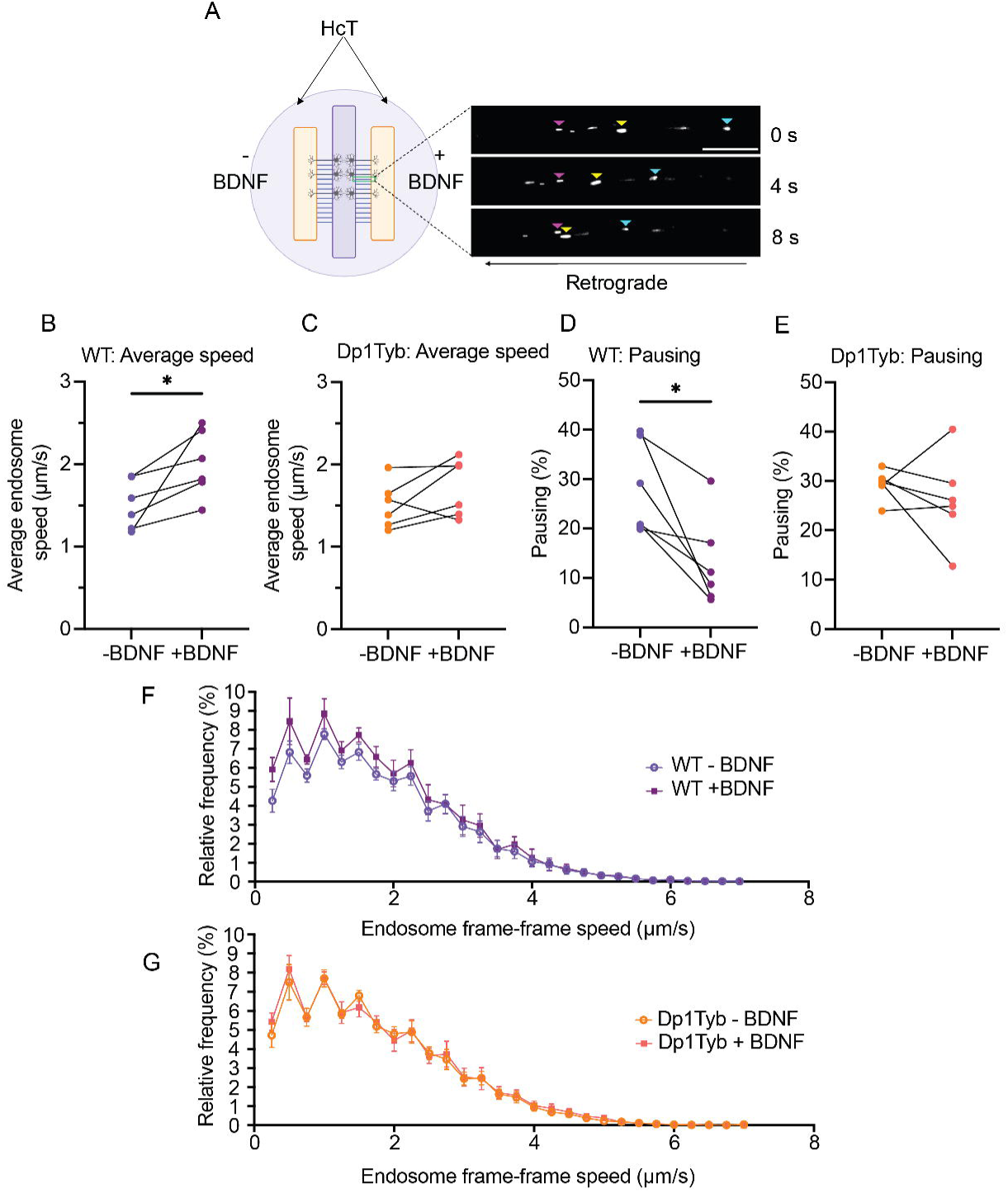
BDNF enhancement of signalling endosome transport is impaired in Dp1Tyb basal forebrain neurons. (A) Schematic of the three-chamber MFC used to culture primary BFNs for axonal transport experiments. Axonal compartments were incubated with Alexafluor555-HcT (HcT-555) ± BDNF and time-lapse imaging was conducted within the MFC microgrooves. Endosomes were manually tracked using TrackMate. Pink arrows highlight a paused endosome, yellow/cyan arrows highlight retrogradely moving endosomes. Scale bar = 20 µm. (B, C) Average endosome speed in WT and Dp1Tyb BFNs in response to BDNF. Lines represent paired conditions within the same three-chambers MFC. BDNF significantly enhances signalling endosome speed in WT (paired Student’s *t* test; *, p=0.0372), but not in Dp1Tyb (paired Student’s *t* test; p=0.1462) BFNs. (D, E) Percentage of pausing of HcT-555-positive signalling endosomes in response to BDNF. BDNF significantly reduces the percentage of pausing in WT (paired Student’s *t* test; *, p=0.0186), but not Dp1Tyb (paired Student’s *t* test; p=0.4451) primary BFNs. (F, G) Speed distribution curves of signalling endosome transport in WT and Dp1Tyb neurons ± BDNF. Endosome pausing (speed <0.25 μm/s) was excluded from the curves). All graphs *N* = 6 biological replicates, 5-8 axons per condition.

Under basal conditions, average signalling endosome transport speeds and percentage of pausing were comparable between WT and Dp1Tyb BFCNs (∼1.5 μm/s and ∼29%, respectively). BDNF significantly increased the average velocities of retrograde signalling endosomes in WT BFNs (∼30%; **Figure 2B, F**) and significantly reduced pausing (>50%; **Figure 2D**). However, in Dp1Tyb BFNs, BDNF treatment did not enhance transport speeds (**Figure 2C, G**) or reduce pausing (**Figure 2E**). Further analysis per axon emphasised these findings (**Supplementary** Figure 2E**, F**).

### Impaired retrograde TrkB trafficking in Dp1Tyb basal forebrain neurons

To assess whether axonal transport abnormalities translate into a deficit in the somatic transfer of internalised TrkB from axon terminals, we used a retrograde accumulation assay previously optimised in the group (19). Twenty-four hours prior to the experiment, the axons of BFNs grown in three-chamber MFCs were incubated with the lipophilic tracers DiI or DiO to identify which neurons had axons projecting into the two axonal compartments. BFNs were then depleted of endogenous growth factors for 2 h prior to supplementing axonal compartments with anti-TrkB antibodies followed by 50 ng/mL BDNF or an anti-BDNF blocking antibody (**Figure 3A**). Immunofluorescence staining of the somatic compartments after 2.5 h revealed TrkB antibodies in the soma of neurons stimulated with BDNF, but not in neurons depleted of BDNF via a blocking antibody (**Figure 3B**). Dp1Tyb BFNs displayed a 30% reduction in the accumulation of TrkB compared to WT BFNs, suggesting reduced axonal retrograde transport of TrkB (**Figure 3C-E**).

**Figure 3:**
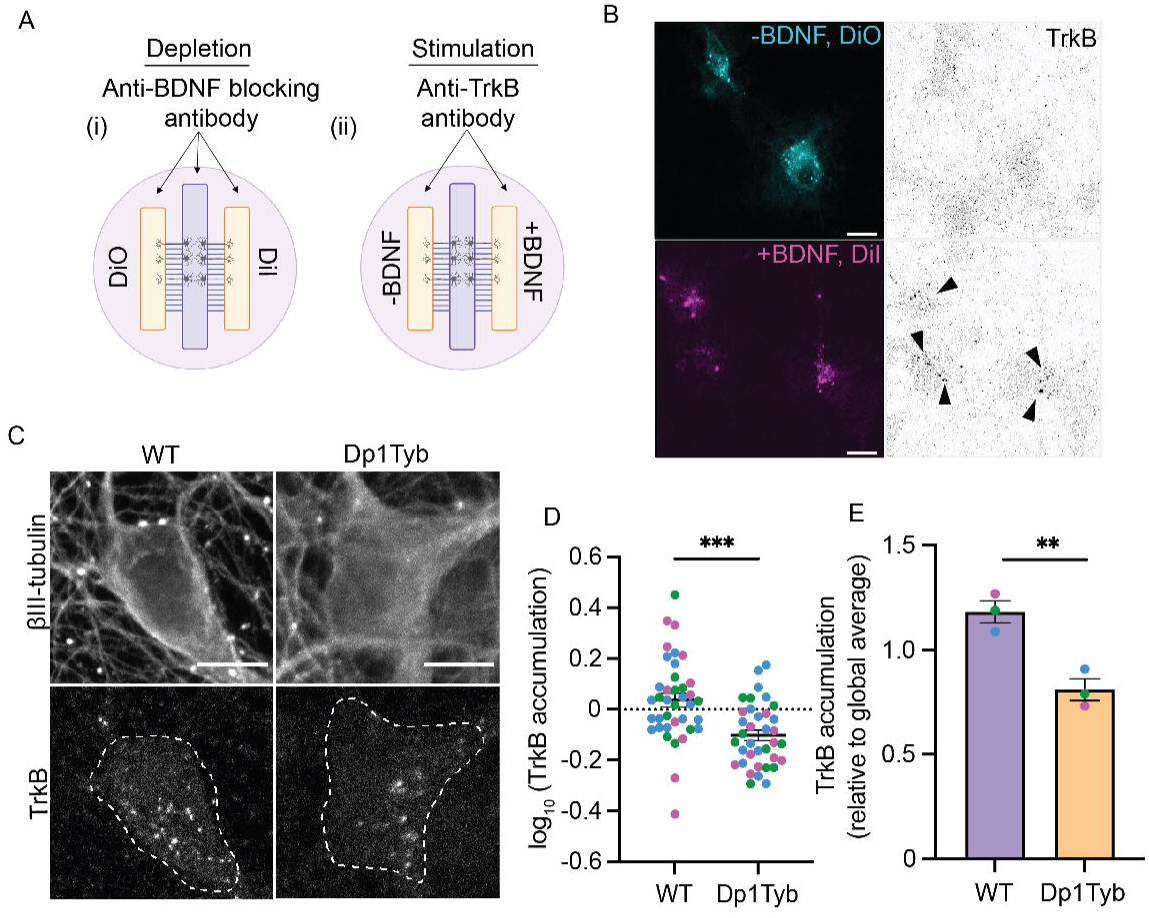
Reduced somatic accumulation of TrkB in Dp1Tyb basal forebrain neurons following axonal stimulation with BDNF. (A) Schematic representation of the TrkB retrograde accumulation assay in three-chamber MFCs. (B) Representative confocal microscopy images of DiI and DiO retrograde tracers, and TrkB accumulation in the somatic compartment ± axonal BDNF treatment. (C) Representative images of TrkB accumulation in the soma following axonal incubation with anti-TrkB antibodies and BDNF in WT and Dp1Tyb neurons. (D, E) A significant decrease in the accumulation of TrkB in Dp1Tyb BFNs per cell (Student’s *t* test; ***, p<0.0001) and per *N* (Student’s *t* test; **, p=0.0073). Different coloured points represent individual *N*s. Bars represent mean ± SEM. *N* = 3 biological replicates. 10-15 cells per biological replicate. Scale bar = 10 µm.

### BDNF signalling is impaired in Dp1Tyb basal forebrain neurons

Next, we wanted to determine whether the signalling pathways downstream of TrkB activation by BDNF were impaired in Dp1Tyb BFNs. Activation of the PI3K-AKT and MAPK-ERK1/2 pathways are crucial for neuronal survival, and alterations in their signalling have been reported in AD-DS models (34). To test these signalling cascades in our system, BFNs were depleted of endogenous growth factors prior to stimulating with 50 ng/mL BDNF for 0, 30, or 120 min and immunostaining for pERK1/2 and pAKT(S473). MAP2 was used as a mask to measure the activation of these signalling cascades within the somatodendritic compartment.

The activation of ERK1/2 appeared significantly impaired in Dp1Tyb BFNs, with a reduction in the amplitude of BDNF response between the two genotypes at 30 min (**Figure 4A, B**). In contrast, this approach did not allow us to detect the effect of BDNF on the activation of pAKT(S473) in either genotype **(Supplementary** Figure 3A**, B**), likely due to high basal signal.

**Figure 4:**
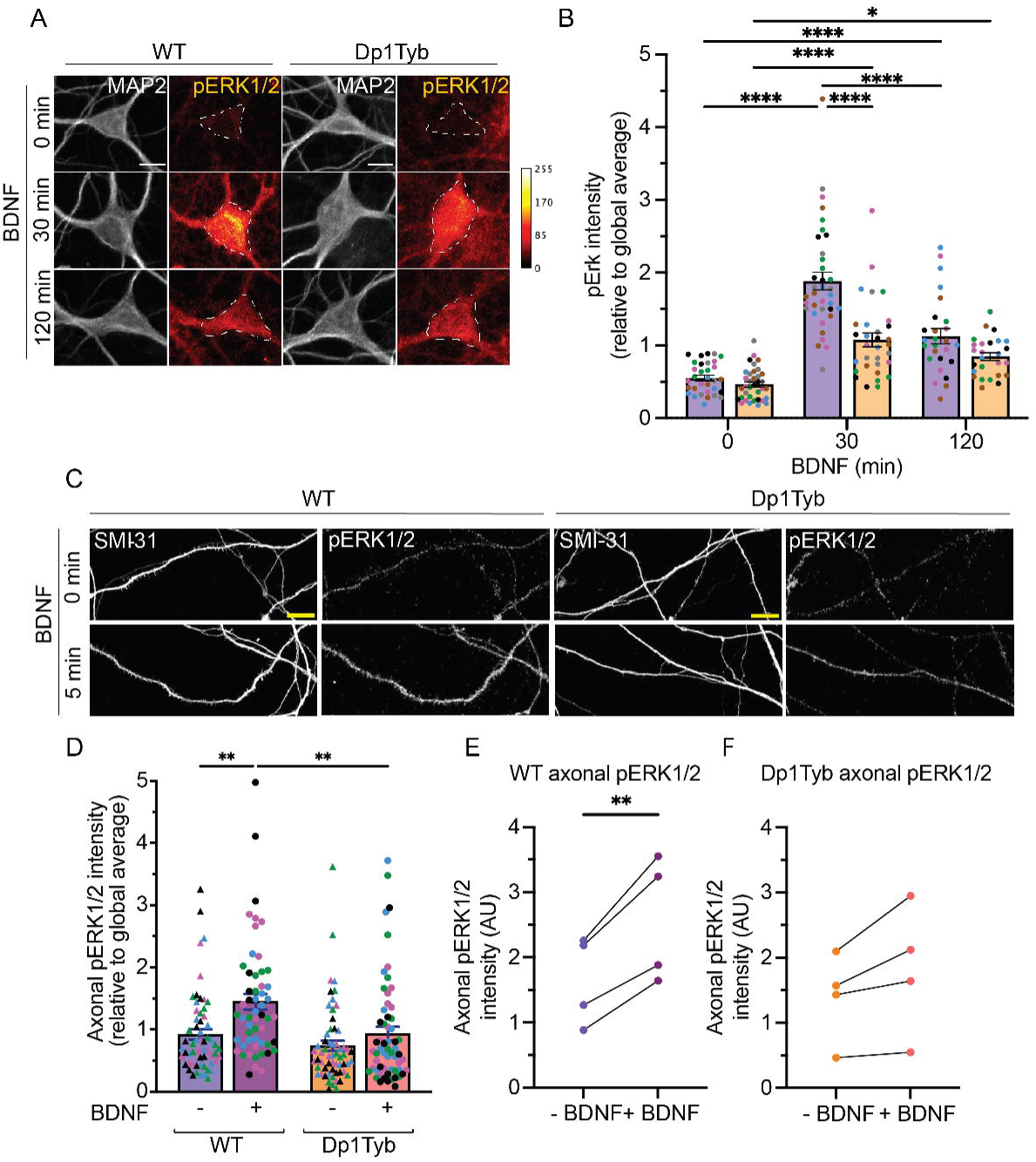
Selective impairment of ERK1/2 signalling following BDNF treatment in Dp1Tyb basal forebrain neurons. (A) Representative confocal images of pERK1/2 staining in WT and Dp1Tyb neurons. (B) Quantification of pERK1/2 immunostaining. There was a significant effect of both BDNF treatment and genotype on ERK1/2 activation (2-way ANOVA with Tukey’s multiple comparisons; BDNF treatment time; ****, p<0.0001, genotype; ****, p<0.0001 and interaction; ****, p<0.0001. Multiple comparisons; ****, p<0.0001, *, p=0.0195). *N* = 5 biological replicates. Bars represent mean ± SEM. Coloured points represent individual *N*s. (C) Representative confocal images of pERK1/2 and SMI-31 staining in WT and Dp1Tyb axons ± BDNF treatment. (D) pERK1/2 intensity per axon in WT and Dp1Tyb BFNs ± 5 min BDNF treatment (2-way ANOVA with Tukey’s multiple comparisons; BDNF treatment time; ***, p=0.0004; genotype; ***, p =0.0006. Multiple comparisons; **, p=0.0024, ****, p <0.0001)). *N* = 4 biological replicates. Bars represent mean ± SEM. Coloured points represent individual *N*s. (E, F) BDNF treatment significantly increased the phosphorylation of ERK1/2 in WT BFN axons (paired Student’s *t* test; **, p=0.0089), but not in Dp1Tyb BFN axons (paired Student’s *t* test; p=0.0915). Scale bar = 10 µm.

Interestingly, these findings do not align with our preliminary data using whole cell lysates probed by western blot (**Supplementary** Figure 3C), which suggested impaired AKT activation in Dp1Tyb cultures (**Supplementary** Figure 3D), and lack of effect of genotype on pERK1/2 levels (**Supplementary** Figure 3E). However, a close inspection of these cultures suggests that the discrepancies in ERK1/2 activation between immunofluorescence and western blot datasets is likely due to differences in the number, and/or activation response of glial cells within the WT and Dp1Tyb BFN cultures. Indeed, we observed increased pERK1/2 staining in non-neuronal cells in Dp1Tyb cultures compared to WT, which further increased upon BDNF treatment, suggesting ERK1/2 activation is TrkB-dependent (**Supplementary** Figure 3F**, G**). These apparently conflicting results highlight the importance of neuron-specific analyses when studying signalling pathways in the context of AD/DS/AD-DS and warrants further investigation.

Given the axonal transport impairment identified above, and the strong link between BDNF-induced ERK1/2 activation and axonal transport (55, 56), we next investigated the activation of ERK1/2 specifically in axons by immunofluorescence. BFNs grown in three-chamber MFCs were depleted of endogenous growth factors prior to treating axons with 50 ng/mL BDNF for 5 min. Whilst treatment for 5 min with BDNF significantly increased the levels of activated ERK1/2 in the axon of WT BFNs (**Figure 4C, D, E)**, there was no significant increase in ERK1/2 activation in Dp1Tyb BFNs (**Figure 4C, D, F**), suggesting impaired axonal ERK1/2 signalling.

### ERK1/2 inhibition in WT neurons mimics the axonal transport impairment identified in Dp1Tyb neurons

To understand whether impaired ERK1/2 activation was contributing to the BDNF-TrkB transport impairment, we treated axons of BFNs cultured in three-chamber MFCs with BDNF ± ERK1/2 inhibitor U0126 (**Figure 5A**). This inhibitor is effective in blocking the activation of ERK1/2 in both WT and Dp1Tyb BFNs as shown in cell lysates by western blot (**Supplementary** Figure 4). We found that impairing ERK1/2 activation by treatment with U0126 caused a 25% reduction in the average axonal transport velocity of signalling endosomes in WT BFNs (**Figure 5B, C, G**). Moreover, U0126 resulted in a 3.4-fold increase in the pausing of signalling endosomes in WT axons (**Figure 5E**), suggesting that the effect of BDNF on the axonal retrograde transport of signalling endosomes is mediated by ERK1/2 activation. In contrast, this ERK1/2 inhibitor had no effect on the axonal retrograde transport of signalling endosomes in Dp1Tyb BFNs **(Figure 5D, F, H)**, reflecting the limited ERK1/2 activation by BDNF in Dp1Tyb BFNs.

**Figure 5:**
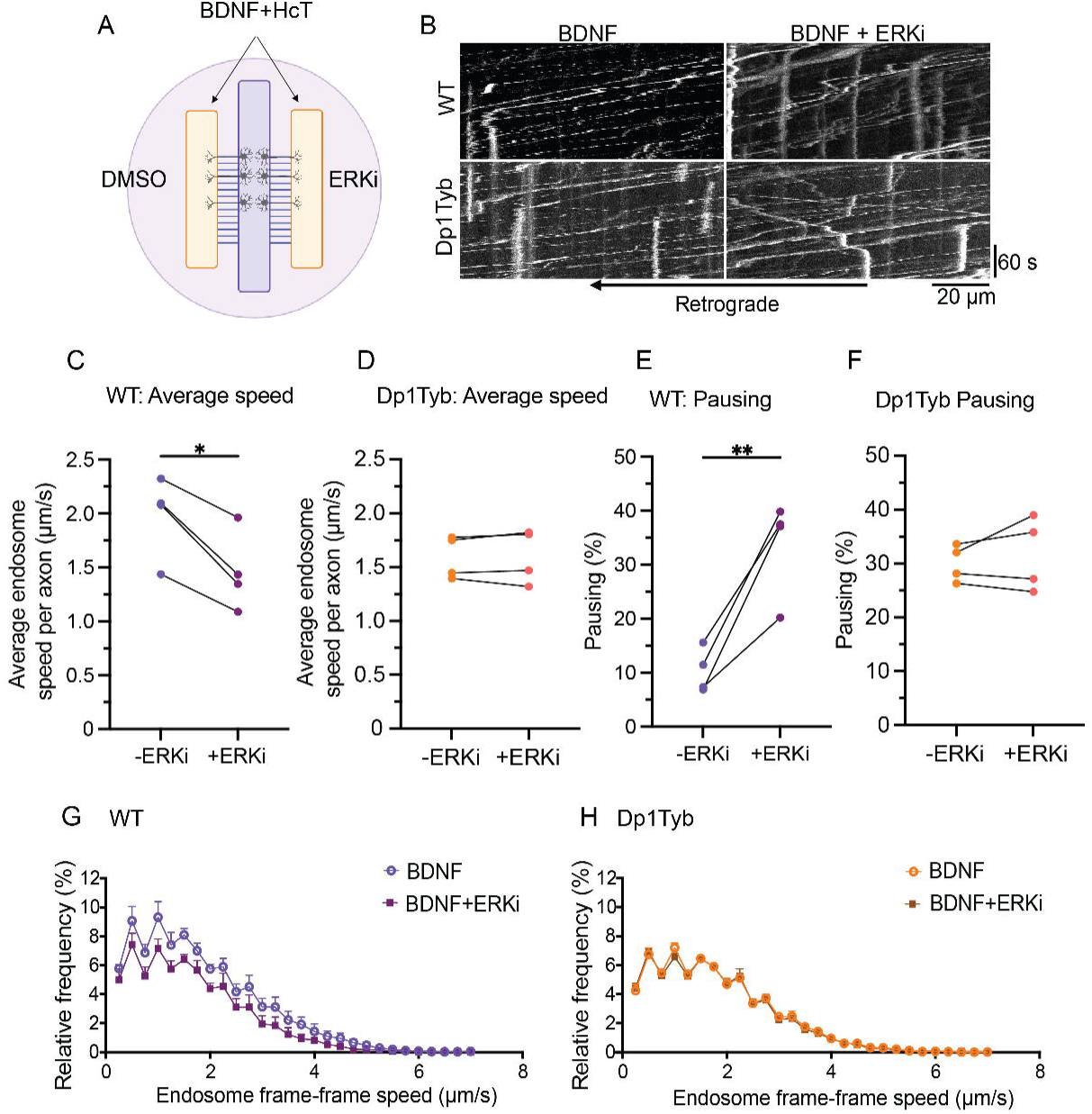
The axonal retrograde transport of signalling endosomes in Dp1Tyb basal forebrain neurons is not affected by ERK1/2 inhibition. (A) Schematic depicting HcT-555 axonal transport experiment ± ERK inhibitor U0126. (B) Representative kymographs of HcT-555-positive signalling endosome transport in WT and Dp1Tyb BFNs ± U0126. (C, D) Average endosome speed per *N* in WT and Dp1Tyb BFNs in response to BDNF ± U0126. Lines represent paired conditions within the same three-chambers MFC. U0126 significantly reduces signalling endosome speed in WT (paired Student’s *t* test; *, p= 0.0135), but not in Dp1Tyb (paired Student’s *t* test; p=0.7136) BFNs. (E, F) Percentage of pausing of HcT-555-positive signalling endosomes in response to U0126 per *N*. Lines represent paired conditions within the same three-chamber MFC. U0126 significantly increases the percentage of pausing in WT (paired Student’s *t* test; **, p=0.0094) but not Dp1Tyb (paired Student’s *t* test; p=0.4571) BFNs. (G-H) Endosome frame-frame speed distribution curve for signalling endosomes in WT and Dp1Tyb primary BFNs ± U0126 treatment. For all graphs *N* = 4 biological replicates, 5-8 axons imaged per condition. Endosome pausing (speed <0.25 μm/s) was excluded from the curves.

## Discussion

Our results identify altered endosomal dynamics and impaired BDNF/TrkB trafficking in Dp1Tyb BFNs, concomitant with decreased ERK1/2 signalling in the axon, a result with potential implications for understanding neurodegenerative processes in AD-DS. We suggest that enlarged early endosomes underpin impaired ERK1/2 signalling, leading to compromised transport of signalling endosomes. Both local and long-distance neurotrophic signalling are crucial for neuronal survival and maintenance. Since physiological growth factor signalling requires functional signalling endosomes, alterations of these organelles due to deficits in sorting, transport and/or homeostatic control, are likely to dampen trophic signalling and affect function both *in vitro* and *in vivo* (32, 43).

Previous BDNF/TrkB transport and signalling studies in AD-DS have been mainly carried out in the Ts65Dn mouse model, which, is confounded by having several extra duplicated mouse genes with an established endocytic function (e.g., *Snx9*, *Synj2*, *Dynlt1*, *Sylt3)* that are unrelated to Chr21 trisomy (49). Furthermore, there is a lack of *in vitro* research investigating the relationship between trafficking and signalling of neurotrophin receptors in BFNs, one of the first populations of neurons to degenerate in AD and AD-DS (27, 57). BFNs are reliant on target-derived trophic support for their survival, thus impaired neurotrophin receptor transport may explain their extreme susceptibility to AD. Based on the shortcomings of the Ts65Dn mouse model, we opted to investigate the post-endocytic trafficking of TrkB in BFNs derived from Dp1Tyb mice. This murine model includes triplication of only those genes on Mmu16 that are syntenic to Hsa21, including the mouse *App* gene. Furthermore, extensive physiological validation of many DS-related phenotypes has been recently provided (47, 48, 58). Here, by utilising a microfluidic system, we uncovered endosomal abnormalities specifically in the axon, directly linking ERK signalling with impaired endosome transport.

Consistent with human post-mortem data in DS (35, 37), we observed a significant enlargement in early endosome size in Dp1Tyb BFNs compared to WT, making our approach relevant for studying endosome dynamics in AD-DS. Elevated Rab5 activity, as shown by increased levels of GTP-bound Rab5, has been suggested to underpin these morphological changes, leading to trafficking defects and neurodegeneration (50). However, data concerning the hyperactivation of Rab5 are often derived from analyses performed in brain tissue, or whole cell lysates, rather than specifically in neurons. In this work, we identified increased levels of GTP-bound Rab5 in Dp1Tyb BFNs. Indeed, the level of active Rab5 significantly increased in WT neurons following 30 minutes of BDNF treatment, whereas it was elevated in Dp1Tyb neurons at steady state.

Recent evidence has shed light onto the pathway involving Rab5 hyperactivation and its downstream effects. Early to late endosome maturation is crucial for maintaining a functional endocytic pathway in neurons. A key step in this maturation process is the switch from Rab5 to Rab7, a transition that is regulated by the Rab7-GEF complex MON1/CCZ1 and requires Rab5 inactivation (59, 60). Hyperactivation of Rab5 may prevent this switch, blocking progression of receptors and other cargo through the endolysosomal pathway, leading to cellular dysfunction. Indeed, in DS, hyperactivation of Rab5 disrupts this equilibrium, leading to failed maturation of endosomal compartments downstream of the early endosome (44). Furthermore, overactivation of Rab5 alone induces an AD-like neurodegenerative cascade both *in vitro* and *in vivo* (50), and antisense oligonucleotides (ASOs) against Rab5 mitigate deficits in neurotrophin signalling, tau phosphorylation and synaptic loss in the Dp16 mouse model of DS (61).

Functional Rab5 is required for the long-distance propagation of the BDNF signal (13). Accordingly, we identified selective deficits in the axonal retrograde transport of BDNF-TrkB signalling endosomes in Dp1Tyb BFNs. Whereas BDNF significantly enhanced signalling endosome speeds and reduced pausing in WT BFNs, as observed in other types of neurons (56, 62–64), these effects were blunted in Dp1Tyb neurons. Rab5 has been associated with endosomes that undergo less processive axonal transport (13, 65). Therefore, if receptors are being retained in enlarged Rab5-positive endosomes, thus preventing their transition to Rab7 organelles, slower velocities and increased pausing might be expected, as we found in Dp1Tyb BFNs. Furthermore, our retrograde accumulation assay demonstrated reduced TrkB transport to the soma, confirming a specific impairment in BDNF-TrkB trafficking in Dp1Tyb BFNs. These findings align with reports of disrupted neurotrophic signalling in models of AD-DS (32, 34, 43).

The binding of BDNF to TrkB triggers the activation of three main signalling pathways: PI3K-AKT, MAPK-ERK1/2 and PLCγ (5). Our data revealed reduced ERK1/2 activation in Dp1Tyb BFNs in response to BDNF, not only in the somatodendritic compartment, but also in axons. Interestingly, dysregulation of early post endocytic trafficking has been suggested to cause signalling impairments. Indeed, disrupting early endosome trafficking through depletion of retrolinkin or endophilin A1 impaired acute ERK activation in culture hippocampal neurons (66). Since ERK1/2 activation is critical for BDNF-mediated endosomal dynamics (55, 56) we hypothesised that inhibiting ERK1/2 might phenocopy the defects observed in the Dp1Tyb BFNs. Consistent with this hypothesis, pharmacologically inhibiting ERK1/2 activation reduced transport speeds and increased pausing in WT BFNs, mirroring the transport deficits seen in Dp1Tyb BFNs. These findings suggest a mutual regulation between trafficking and signalling, wherein defects in sorting from early endosomes impair ERK1/2 activation, which further disrupts axonal transport. This dynamic interplay highlights a bidirectional relationship between endosomal signalling and trafficking, potentially amplifying pathological effects.

ERK1/2 activation has been shown to enable the recruitment of the dynein intermediate chain to signalling endosomes in neurons (55). Furthermore, expression of an intermediate chain that could not be phosphorylated by ERK1/2 and recruit TrkB resulted in a 45% decrease in cell survival, highlighting the importance of ERK1/2-dependent signalling for long-distance neurotrophin functions (55). Crucially, we found that ERK1/2 inhibition had no effect on axonal transport in Dp1Tyb neurons. The lack of effect is consistent with an impairment of ERK1/2 in Dp1Tyb neurons occurring downstream of early endosome abnormalities. These upstream defects likely limit the capacity of BDNF to elicit a physiological signalling response, making the pharmacological inhibition of ERK1/2 inconsequential to the already compromised transport and signalling dynamics.

Retrograde transport relies on cytoplasmic dynein forming a complex with its cofactor, dynactin, and a range of dynein adaptors (67). ERK1/2 activation might result in an increased local translation of dynein cofactors and/or adaptor proteins. Indeed, mRNAs encoding dynein regulators have been found in the axon in the central and peripheral nervous systems (68–70). In dorsal root ganglia (DRG) neurons, axonal transcripts of the dynein adaptor protein LIS1, and the dynactin subunit p150^Glued^ are translated in response to increased NGF signalling. Inhibition of protein synthesis using anisomycin, or knockdown of mRNA transcripts for *LIS1* or *p150*^Glued^, abolished the NGF-induced increase in the retrograde transport of lysosomes and quantum dots-conjugated NGF (70). Interestingly, only NGF-induced, rather than basal, axonal transport seems dependent on local translation, possibly reflecting previously inactive dynein complexes becoming activated and bound to cargo in response to NGF. This result might be relevant to the present study given that the transport deficits in Dp1Tyb BFNs are only evident upon BDNF stimulation, suggesting that neurotrophin signalling endosomes are specifically impaired.

Similar mechanisms have been identified in primary motor neurons, with IGFR1 inhibition leading to increased local translation of the dynein adaptor BICD2 and the subsequent enhancement of signalling endosome transport (53, 62). In addition, BDNF signalling drives the phosphorylation of several microtubule associated proteins (MAPs) in human iPSC-derived lower motor neurons (64), and ERKs have been shown to regulate the microtubule landscape (71). The impairment upon BDNF stimulation in Dp1Tyb BFNs may therefore reflect defects in local protein translation of dynein associated proteins in axons, or altered MAP phosphorylation, due to altered ERK1/2 activation.

The dysregulation of early endosome morphology and function, coupled with impaired BDNF transport and signalling, might highlight a potential convergence of pathogenic mechanisms in AD-DS. Our results emphasize the need for targeted therapeutic strategies that address endosomal trafficking defects and signalling impairments. Future studies should investigate the precise molecular mechanisms linking increased levels of GTP-Rab5 and dampened ERK1/2 signalling to TrkB transport deficits in AD-DS models.

## Materials and methods

### Animals

C57BL/6J;129P2-Dp(16*Lipi*-*Zbtb21*)1TybEmcf mice (hereafter referred to as Dp1Tyb) were maintained on a C57BL/6J background for more than ten generations (47, 48). Animals were housed in a temperature and humidity-controlled environment with a 12 h light/dark cycle, with access to food and water ad libitum. Timed matings were set with Dp1Tyb males and C57BL/6J females. Embryos were collected 16 days after the detection of a positive plug (E16) and genotype confirmed by PCR using the following primer pairs (Forward: CTAACCCTAACCCTAAGTCCTTGTC, Reverse: TGAGGAGAGTTTTCTGGGAGAA; Forward: CGGGCCTCTTCGCTATTACG, Reverse: GAGTGCATTCTCCTCATTCAGATG) and PCR cycling conditions (initial denaturation at 94°C for 3 min; 35 cycles of denaturation at 94°C for 30 s, annealing at 57°C for 10 s, and elongation at 72°C for 1 min; followed by a final extension at 72°C for 7 min).

All experiments were conducted under the guidelines of the UCL Genetic Manipulation and Ethics Committees and in accordance with the European Community Council Directive of 22 September 2010 (2010/63/EU). All procedures for the care and treatment of animals were carried out under license from the U.K. Home Office in accordance with the Animals (Scientific Procedures) Act 1986 Amendment Regulations 2012.

### Plasmids and Reagents

Reagents were from Sigma-Aldrich unless otherwise stated. Recombinant human NGF and BDNF were purchased from PeproTech. pGEX-4T-2/Rabaptin-5:R5BD was a gift from Guangpu Li (Addgene plasmid # 168808; http://n2t.net/addgene:168808; RRID:Addgene_168808) (72). ERK inhibitor U1026 was from Promega (V1121). Primary antibodies for immunofluorescence (IF) and western blot (WB) were sourced as follows: MAP2 (Abcam, ab5392, RRID:AB_2138153, IF 1:2,000), SMI-31 (BioLegend, 801601, RRID:AB_2564641, IF 1:300), ERK1/2 (4695, RRID:AB_390779, IF 1:250, WB 1:1,000), pERK1/2_Thr202/Tyr204_ (9101, RRID:AB_331646, IF 1:250, WB 1:1,000), AKT (9272S, RRID:AB_329827, IF 1:250, WB 1:1,00), pAKT(Ser473) (Cell Signalling Technology 4060S, RRID:AB_2315049, IF 1:250, WB 1:1,000), TrkB (R&D Systems, AF1494, RRID:AB_2155264, IF 1:250, WB 1:500), EEA1 (BD Transduction Laboratories, 610457, RRID:AB_397830, IF 1:500), GST (Abcam, ab9085, RRID:AB_306993, IF 1:500), and GADPH (Sigma Aldrich, MAB374, RRID:AB_2107445, WB 1:1,000). Antibodies used for the retrograde accumulation assay were as follows: TrkB (Merck (Millipore) AB9872, RRID:AB_2236301, 1:50) and BDNF (BioTechne, AF248, RRID:AB_355275, 1:50).

### BFN culture

The basal forebrain (BF) from Dp1Tyb and WT embryos of either sex was dissected at E16 in Hank’s balanced salt solution (HBSS) and transferred in Hibernate E medium (ThermoFisher, #A1247601). Tissue was washed in HBSS prior to incubating in Accumax (Innovative Cell Technologies, #AM105) for 10 min, then diluted (1:1) in HBSS for a further 5 min. Tissue was resuspended in warm plating medium (Minimum Essential Medium (MEM; Gibco) supplemented with 10% heat inactivated horse serum, 38 mM NaCl, 0.6% glucose and 1 × GlutaMAX (Invitrogen) prior to mechanical dissociation by pipetting. 10-15,000 cells per cm^2^ were seeded onto glass coverslips or microfluidic chambers (MFCs) pre-coated with 1 mg/mL poly-L-lysine. Fabrication and coating of MFCs was conducted as described previously (62). Two hours after plating, medium was changed to maintenance medium (Neurobasal; Gibco) supplemented with 1% penicillin/streptomycin (ThermoFisher), 38 mM NaCl, 0.6% glucose, 1× GlutaMAX, 1× B27 (Gibco), 10 ng/mL NGF and 10 ng/mL BDNF). Cultures were maintained in a humidified incubator at 37 °C, 5% CO_2_. Medium was half changed every 3-5 days. After 3 days *in vitro* (DIV) the axonal compartment of each MFC was supplemented with 20 ng/mL BDNF and NGF to encourage axonal growth; the somatic compartment was maintained with 10 ng/mL BDNF and NGF.

### Recombinant proteins

The Rabaptin-5(R5BD) fusion protein was produced in chemically competent NEB 5-alpha *E. coli* (New England Biolabs) transformed with the pGEX-4T-2/Rabaptin-5:R5BD plasmid. Protein expression was induced by treatment with 0.5 mM isopropyl β-D-1-thiogalactopyranoside (IPTG) for 4 h. Cells were harvested by centrifugation at 4,000 × g for 30 min at 4 °C. The pellet was resuspended in 50 mM Tris HCl (pH 7.5), 150 mM NaCl, 0.05% Tween-20, 2 mM EDTA, 0.1% 2-mercaptoethanol (βME) and Halt protease and phosphatase inhibitor cocktail (1:100, ThermoFisher). Bacteria were lysed by sonication with 4 x 15 s pulses at 40 kHz. The bacterial lysate was centrifuged at 10,000 × *g* for 40 min and the pellet discarded. Rabaptin-5(R5BD) was purified using glutathione-agarose beads (Merck-Millipore) as per manufacturer’s instructions and eluted using 10 mM reduced glutathione (pH 8.0). Purified proteins were dialysed using Slide-A-Lyzer G2 Dialysis Cassettes (ThermoFisher, #87735) to remove the reduced glutathione and Ultra-0.5 mL centrifugal filters with a 10 kDa cut off (Amicon) were used to concentrate proteins post dialysis.

### Rab5 activity assay

Rab5 activity was determined using a protocol adapted from Moya-Alvarado *et al*., 2018 (17). Neurons cultured on coverslips were depleted of endogenous growth factors for 2 h prior to stimulation with 50 ng/mL BDNF for 0, 5 or 30 min. Cells were washed in PBS and briefly fixed for 8 min with 4% PFA, 4% sucrose in PBS before blocking for 45 min in 3% BSA reconstituted in incubation buffer (50 mM Tris-Cl, 50 mM NaCl, 5 mM MgCl_2_, 1 mM EDTA, 0.25 mM sucrose, 0.2% Triton X-100 and 0.5 mM dithiothreitol (DTT) added fresh; pH 7.2). GST-Rabaptin5R5BD and GST were purified and diluted to 15 μg/mL in incubation buffer. 10 μM reduced glutathione was added to the buffer to reduce formation of aggregates. Diluted proteins were centrifuged at 20,000 × g for 30 min prior to applying to the neurons overnight at 4 °C with shaking. The following day cells were washed briefly 5 times with PBS and refixed for 15 min with 4% PFA, 4% sucrose for anti-GST immunofluorescence.

### Axonal retrograde transport

At DIV 14, neurons grown in three-chamber MFCs were depleted of endogenous growth factors for 2 h in Neurobasal. For the final 30 min of depletion, axonal compartments were supplemented with 30 nM AlexaFluor555-H_c_T (53, 62). Axons were incubated in the presence or absence of 50 ng/mL BDNF for 30 min. All media was then replaced with Neurobasal supplemented with 20 nM HEPES-NaOH (pH 7.4) and imaged (Zeiss LSM 980 with Airyscan 2). In the BDNF conditions, the Neurobasal/HEPES-NaOH was supplemented with 50 ng/mL BDNF in the axonal compartment for imaging. During image acquisition, neurons were kept at 37 °C in the humidified chamber of the microscope. Each MFC was imaged for approximately 40 min, acquiring videos at 2 frames/s from 5-8 axons per condition. For ERK1/2 inhibitor experiments, axonal compartments were incubated with 50 ng/mL BDNF, 30 nM H_c_T in the presence of 20 μM U0126 or an equivalent amount of DMSO.

### Somatic accumulation of TrkB

The retrograde accumulation of TrkB was assessed using an established protocol with minor modifications (19). Neurons were grown in three-chamber MFCs. 24 h before the experiment, axonal compartments were incubated with 10 ng/mL of the lipophilic retrograde tracers DiI (Invitrogen, D3911) and DiO (Invitrogen, D275). Having two different retrograde tracers enables identification of neurons projecting axons to each compartment. On DIV14, the media was replaced with Neurobasal and an anti-BDNF blocking antibody (1:50, BioTechne, AF248). After 2 h, axonal compartments were washed with Neurobasal and axons incubated with polyclonal antibodies against the extracellular domain of TrkB (Merck Millipore, AB9872) together with 50 ng/mL BDNF or an anti-BDNF blocking antibody (1:50). After 2.5 h, neurons were fixed with 4% PFA, 4% sucrose in PBS. To reveal the somatic accumulation of TrkB antibodies, cells were blocked and permeabilised in 4% BSA, 0.2% saponin for 1 h followed by incubation with Alexa Fluor-conjugated secondary antibodies.

### Signalling assays

Neurons cultured on coverslips, in 6-well plates, or MFCs were depleted of endogenous growth factors by incubation in Neurobasal for 2 h before stimulation with 50 ng/mL BDNF for 0, 5, 30 or 120 min to establish a time course of ERK1/2 and AKT activation in BFNs. Cells on coverslips and MFCs were fixed with 4% PFA, 4% sucrose, and the phosphorylation of ERK1/2 and AKT in the soma and/or axons was analysed by IF and confocal microscopy. Following the BDNF time course in 6-well plates, cells were lysed on ice in RIPA buffer (50 mM Tris-HCl pH 7.5, 150 mM NaCl, 1% NP-40, 0.5% sodium deoxycholate, 0.1% sodium dodecyl sulphate, 1 mM EDTA, 1 mM EGTA) with Halt protease and phosphatase inhibitor cocktail (1:100, ThermoFisher), and the phosphorylation of ERK1/2 and AKT was analysed by western blotting.

### Immunofluorescence

Neurons were washed in PBS and fixed for 15 min in 4% PFA, 4% sucrose prior to incubation in 0.15 M glycine (pH 7.0) in PBS for 10 min. Neurons were blocked and permeabilised in 4% BSA, 0.2% saponin in PBS for 1 h before incubating with primary antibodies diluted in 4% BSA, 0.05% saponin in PBS overnight at 4 °C. Samples were washed three times with PBS and incubated for 1 h with AlexaFluor-conjugated secondary antibodies and 4L,6-diamidino-2-phenylindole (DAPI) (1:500) diluted in 4% BSA, 0.05% saponin in PBS. Finally, neurons were washed in PBS and mounted using Fluoromount-G (Invitrogen).

### Western blotting

Purified proteins or cell lysates were mixed with 4× Laemmli buffer (250 mM Tris-HCl, pH 6.8 8% SDS, 40% glycerol, 0.02% bromophenol blue) and 10% fresh βME and denatured at 100 °C for 5 min. BCA protein Assay Kit (ThermoFisher) was used to quantify the protein concentration. Proteins were separated using 4–15% Mini-PROTEAN TGX Stain-Free Protein Gels (Bio-Rad) and transferred onto methanol-activated polyvinylidene difluoride (PVDF) membranes with the semi dry Trans-Blot Turbo Transfer system (Bio-Rad). Membranes were blocked in 5% BSA in PBS with 0.1% Tween 20 (PBS-T) for 1 h at room temperature. Primary antibodies were incubated overnight at 4 °C with shaking. Membranes were washed three times with PBS-T for 10 min before incubation with the appropriate horseradish peroxidase (HRP) conjugated secondary antibody (DAKO) for 1 h at room temperature. After three 10 min washes with PBS-T, Immobilon Crescendo Western HRP Substrate (Merck-Millipore) was used for chemiluminescent detection of proteins using the ChemiDoc Touch Imaging System.

### Image acquisition, quantification, and statistical analysis

All images were acquired using the Zeiss LSM 980 confocal microscope with Airyscan 2 using an oil immersion 40×/1.3 numerical aperture (NA) or 63×/1.4 NA objectives unless otherwise stated. Fixed samples were imaged as Z-stacks. For time-lapse imaging of live neurons, images were acquired at 2 frames/s. Images were analysed using Image J/FIJI (version 2.1.0, 1.53c) (73). For the analysis of signalling endosome dynamics, the FIJI plugin Trackmate (74) was used to manually track all retrogradely moving organelles. Endosomes were selected and manually connected in each adjacent frame giving single frame-frame velocities which were then averaged across the entire track, giving an average speed for each endosome. The endosome speed was then averaged per axon and per biological replicate (N). A biological replicate is defined as mouse embryos of the same genotype pooled from one litter. Speed distribution curves were generated to display the relative frame-frame movement of all signalling endosomes. A pause was defined as a signalling endosome that moved with a speed less than 0.25 μm/s between consecutive frames. The percentage of time paused was determined by the number of pauses divided by the total number of frame-frame movements per axon. This was then averaged per *N*.

Western blot bands were quantified using Image Lab (Bio-Rad, version 6.1.0).

### Statistical analysis

Data was analysed and visualised using GraphPad Prism 9 (La Jolla, California, USA, version 9.4.1) and tested for normality and homoscedasticity. Where assumptions were not met, data was log transformed. For analyses with >2 groups, a two-way ANOVA was used to compare means followed by Tukey’s multiple comparisons. For comparison of two groups, Student’s *t-test* was used. Levels of significance are indicated in figure legends. Mean ± standard error is shown.

## Acknowledgements

We thank the personnel of the MRC Prion Unit BSF for assistance in maintaining the mouse colonies. We thank Dr Sunaina Surana for help with purifying GST-tagged proteins.

## Author Contributions

Conceptualisation: EB, OML, GS. Investigation: EB. Writing: Original Draft, EB; Editing: EB, NB, EMCF, OML, GS. Supervision: NB, EMCF, GS. Funding: EB, GS.

## Ethics Statement

All experiments were conducted under the guidelines of the UCL Genetic Manipulation and Ethics Committees and in accordance with the European Community Council Directive of 22 September 2010 (2010/63/EU).

## Data Availability

All data are provided in the main text.

