## Supplementary figures and images for "Impaired BDNF-TrkB trafficking and signalling in Down syndrome basal forebrain neurons"

### Supplementary Figure 1

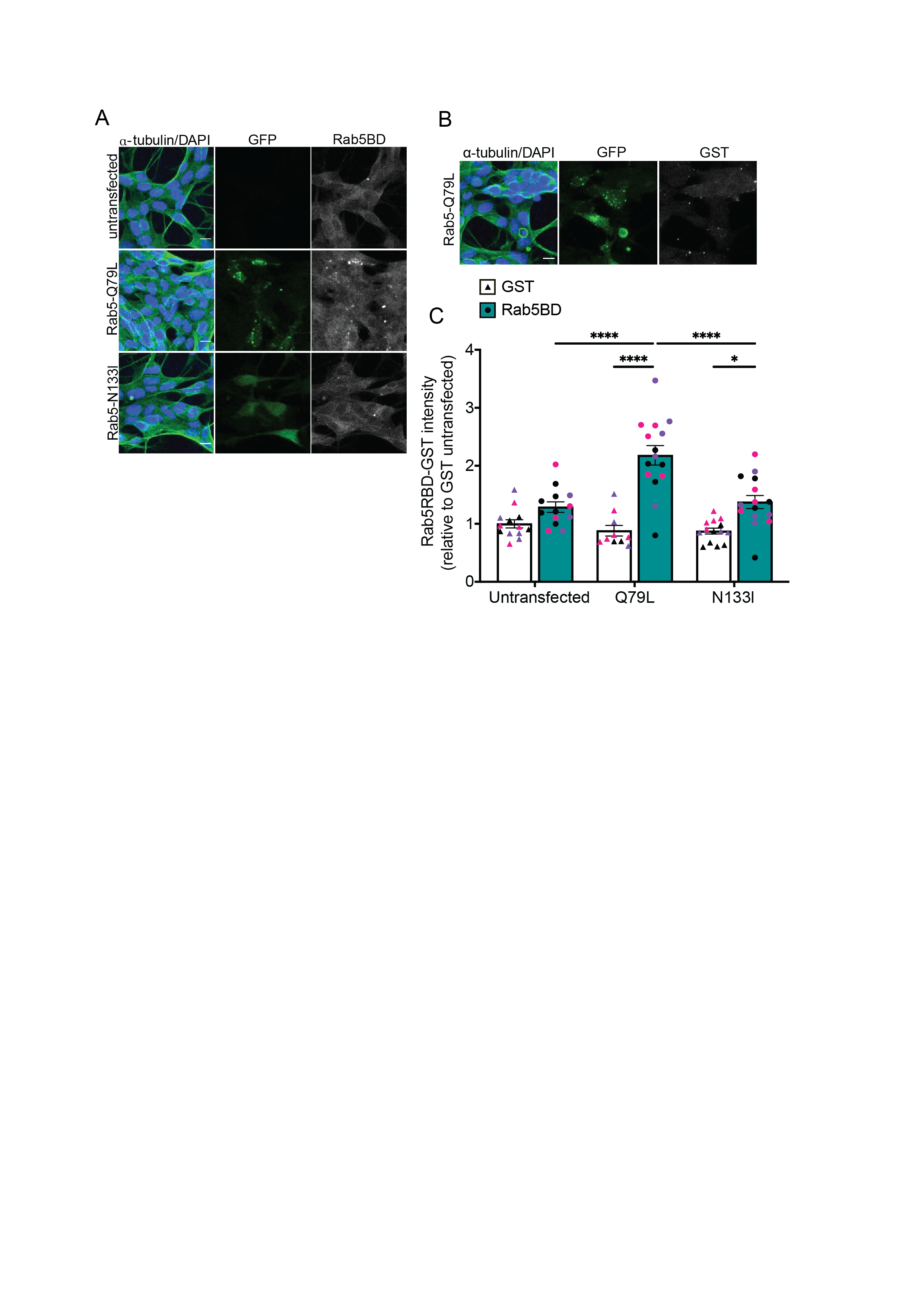

### Supplementary Figure 2

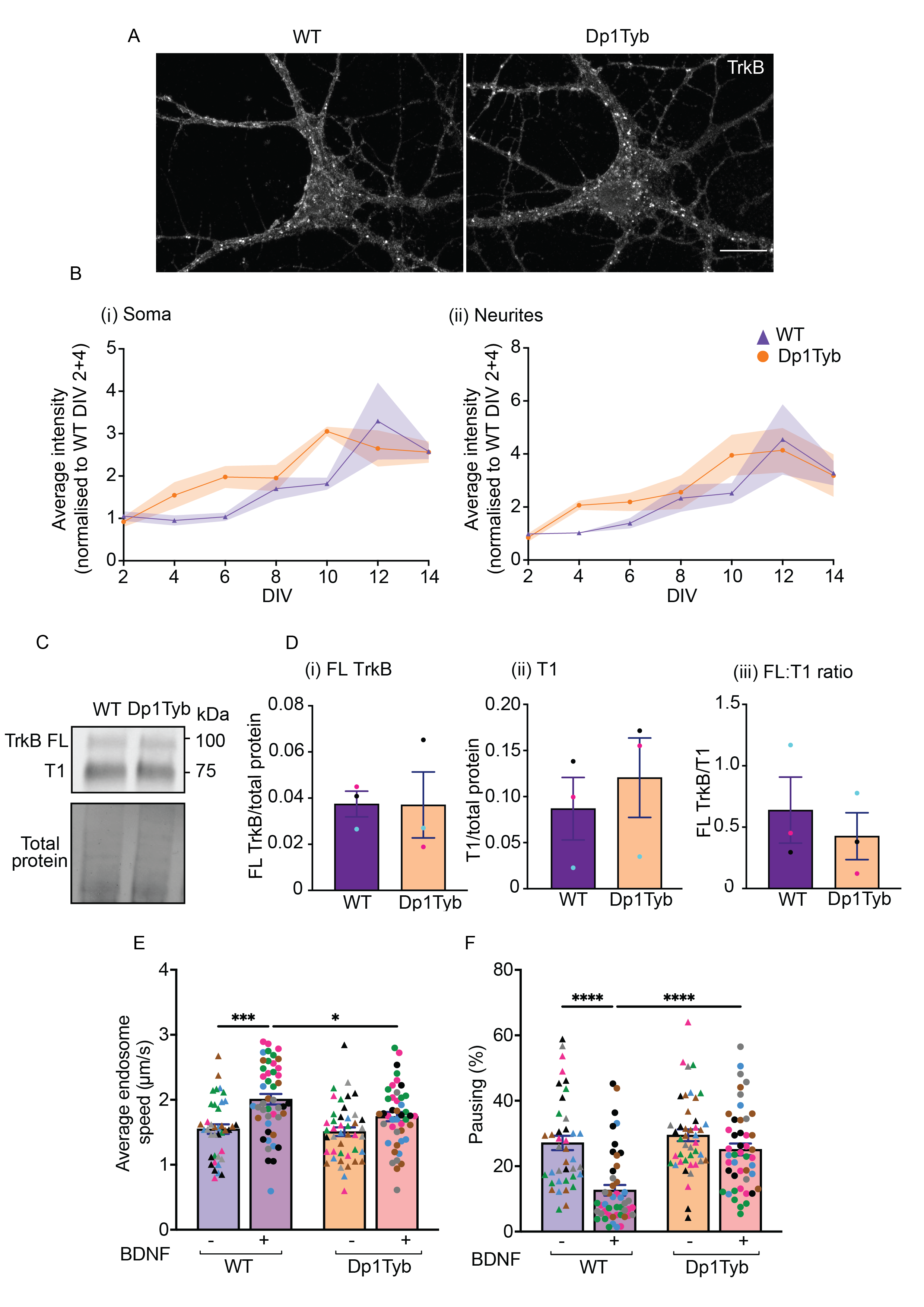

### Supplementary Figure 3

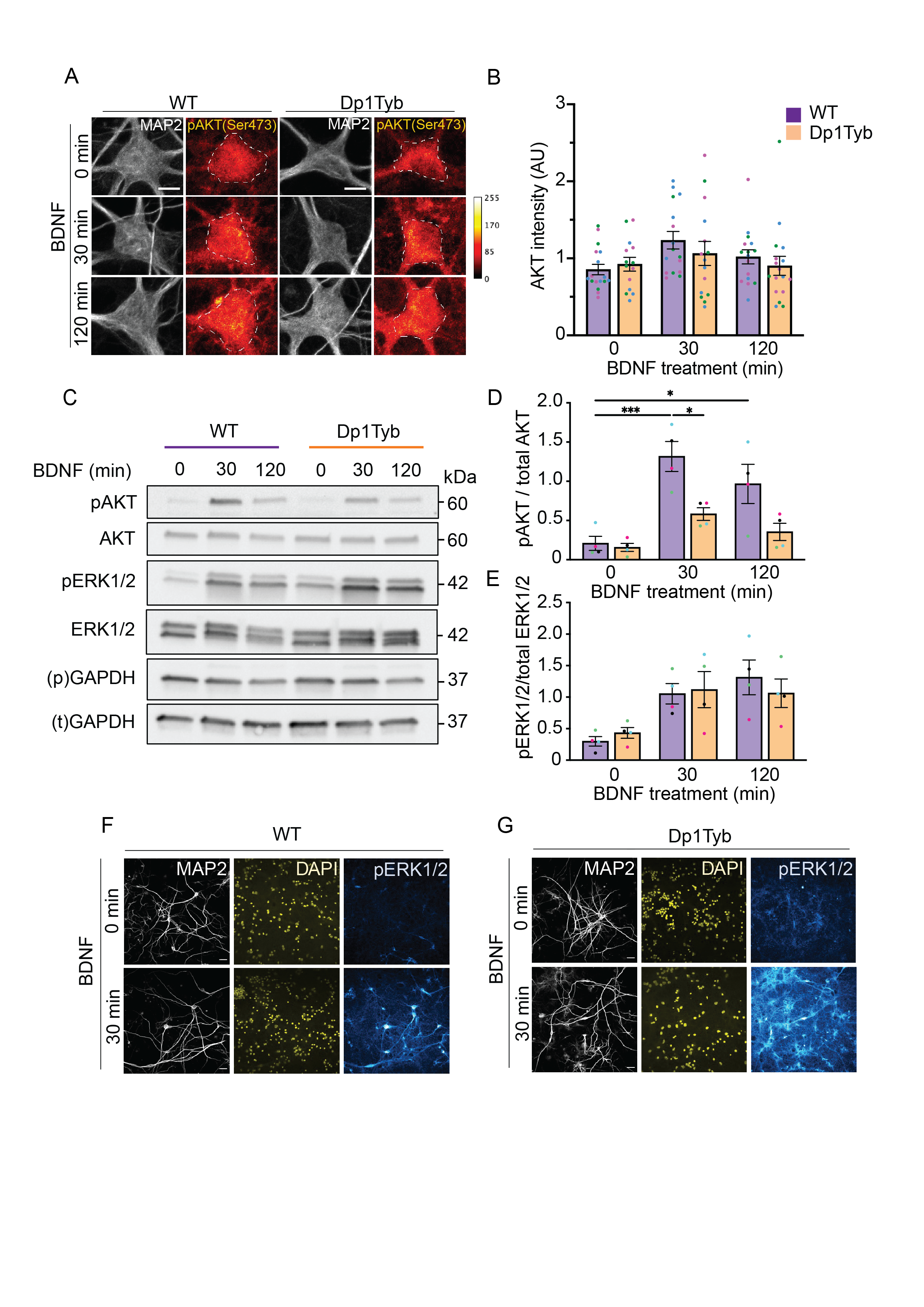

### Supplementary Figure 4

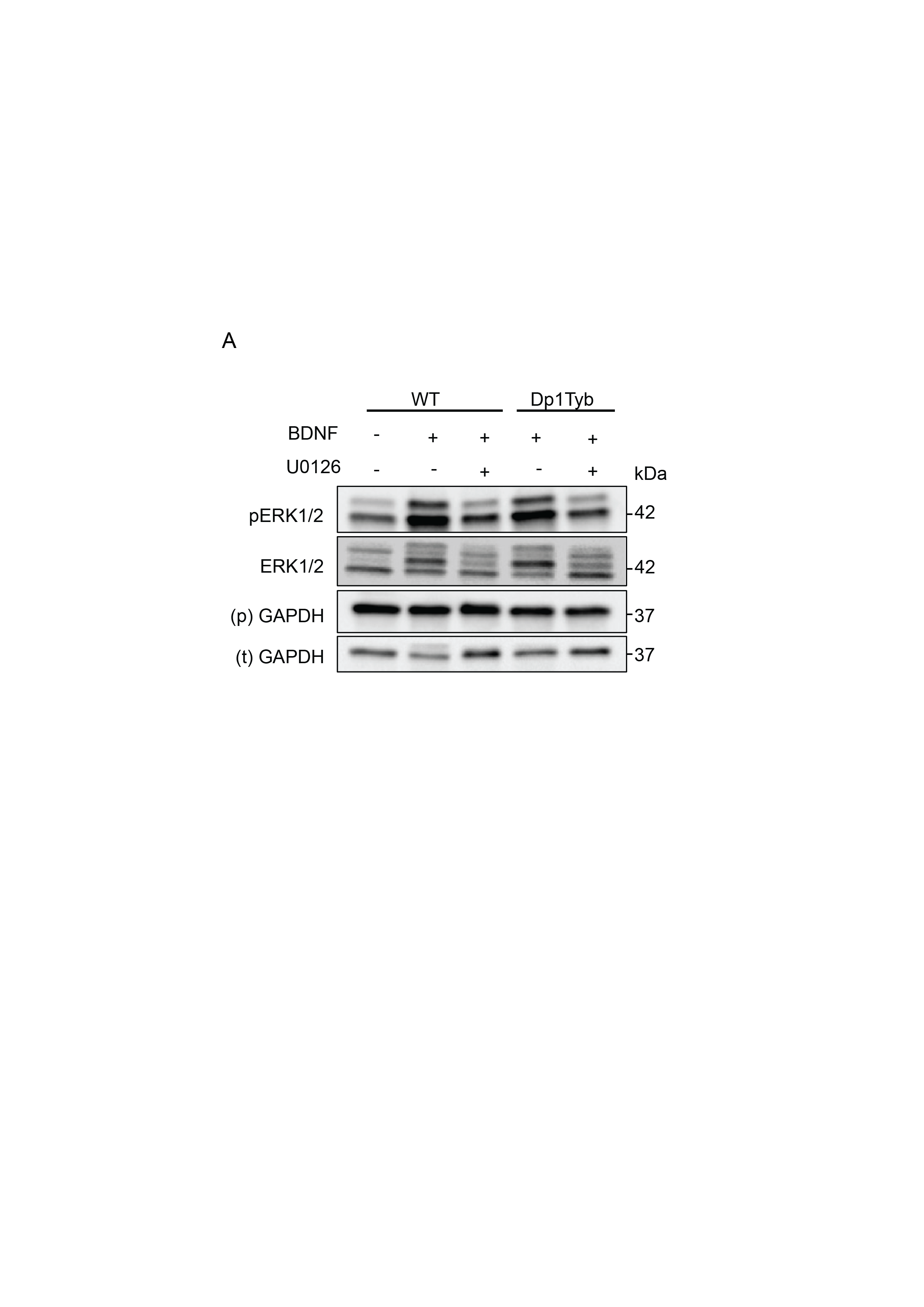
